# Heteroplasmy is rare in plant mitochondria compared to plastids despite similar mutation rates

**DOI:** 10.1101/2023.11.15.567200

**Authors:** Marina Khachaturyan, Mario Santer, Thorsten B. H. Reusch, Tal Dagan

**Affiliations:** Marine Evolutionary Ecology, GEOMAR Helmholtz Centre for Ocean Research Kiel, Kiel, Germany; Institute of General Microbiology, University of Kiel, Kiel, Germany; Department of Human Molecular Genetics & Biochemistry, Faculty of Medicine, Tel Aviv University, Tel Aviv, Israel

**Keywords:** plant organelle evolution, plastids, mitochondria, eelgrass, substitution rate, genetic diversity, allele dynamics, heteroplasmy, simulated evolution, *Zostera marina*

## Abstract

Plant cells harbor two membrane-bound organelles containing their own genetic material – plastids and mitochondria. Although the two organelles co-exist and co-evolve within the same plant cells, they differ in genome copy number, intracellular organization, and mode of inheritance. How these attributes determine the time to fixation, or conversely, loss of neutral alleles is currently unresolved. Here we show that mitochondria and plastids share the same mutation rate yet plastid alleles remain in a heteroplasmic state significantly longer compared to mitochondrial alleles. By analysing genetic variants across populations of the marine flowering plant *Zostera marin*a and simulating organelle allele dynamics, we examine the determinants of allele segregation and fixation time. Our results suggest that bottleneck on the cell population, e.g., during branching and seeding, and stratification of the meristematic tissue, are important determinants of mitochondrial allele dynamics. Furthermore, we suggest that the prolonged plastid allele dynamics are due to a yet unknown active plastid partition mechanism. The dissimilarity between plastid and mitochondrial novel allele fixation at different levels of organization may figure into differences in adaptation processes. Our study uncovers fundamental principles of organelle population genetics that are essential for further investigations of long-term evolution and molecular dating of divergence events.

## Introduction

Genetics of the eukaryotic organelles – plastid and mitochondria – share multiple characteristics with the genetics of other extrachromosomal genetic elements. For example, the organelle DNA (oDNA) copy number may significantly outnumber that of the nuclear chromosomes, similar to prokaryotic plasmids (reviewed in Rodríguez-Beltrán et al. 2021). The high copy number entails the possibility of intra-cellular genetic diversity, termed ‘heteroplasmy’ (reviewed in Ramsey and Mandel 2019). The intermediate intracellular allele frequency changes due to random allele segregation during the organelle and cell division, termed ‘segregational drift’ (Birky et al. 1989). Segregational drift of plastid alleles occurs independently during plastid and cell division. The effect of mitochondria division on allele segregation is considered negligible due to regular mitochondrial fusion and fission (Lonsdale et al. 1988; Logan 2010; Rose 2019). Plastids and mitochondria are typically considered to be uniparentally inherited, in contrast to the biparentally inherited nuclear genome, although organelle paternal (and maternal) leakage was previously reported (reviewed in Ramsey and Mandel 2019). Assuming strict uniparental inheritance, organelles evolve analogous to asexual microbes, with new variants arising solely via *de novo* mutations. Consequently, organelle genomes are expected to accumulate slightly deleterious mutations over time, eventually leading to mutational meltdown (Gabriel et al. 1993). The main factor that mitigates mutational meltdown in plant organelles is thought to be their lower mutation rate compared to the nuclear genome (Palmer and Herbon 1988; Lynch et al. 2006). Additionally, a high level of interchromosomal recombination (gene conversion) was suggested to prevent the organelle mutational meltdown (Khakhlova and Bock 2006; Maréchal and Brisson 2010; Edwards et al. 2021).

The maintenance of organelles upon cell division requires their even partitioning into daughter cells. Prior to cell cytokinesis, mitochondria fission into small entities that are unbiasedly scattered in the cytoplasm (Sheahan et al. 2004). The number of small mitochondria may exceed the mitochondrial DNA (mtDNA) copy number (Arimura et al. 2004; Preuten et al. 2010). An even cytoplasm volume division between the two daughter cells thereby ensures that the mtDNA distribution is balanced (Sheahan et al. 2004; Sheahan et al. 2007). In contrast, plastids are localized in the perinuclear area prior to the cell division, where the number of plastids is doubled by plastid division. During cytokinesis, plastids cluster around the daughter nuclei, which may ensure their segregation. The proper plastid positioning throughout the cell cycle is ensured by actin filaments (Sheahan et al. 2004). However, how plastids are evenly partitioned into the daughter cells remains unclear.

In multicellular organisms, the accumulation of mutations in nuclear and organelle genomes is slowed down within the germline. Recent studies suggest that few slowly dividing pluripotent stem cells in the meristem central zone – termed ‘shoot apical initials’ and referred to as ‘stem cells’ – later form a transient sexually active plant germline (Figure 1; Kwiatkowska 2008; Lanfear 2018; Burian 2021). The stem cell population undergoes regular bottlenecks during stem branching and seed formation. Stem cells divide preferentially within the corresponding meristem layer, which comprises 1-2 tunica layers and an internal corpus layer. Most cell divisions of stem cells are asymmetrical, i.e., only one daughter cell remains a stem cell. Symmetrical cell division yields two daughter stem cells, where cell differentiation eventually preserves the total number of initials on the apex (Figure 1; Bowman and Eshed 2000; Morrison and Kimble 2006; Burian 2021). The effect of the plant meristem structure on the oDNA allele dynamics remains understudied.

**Fig. 1:**
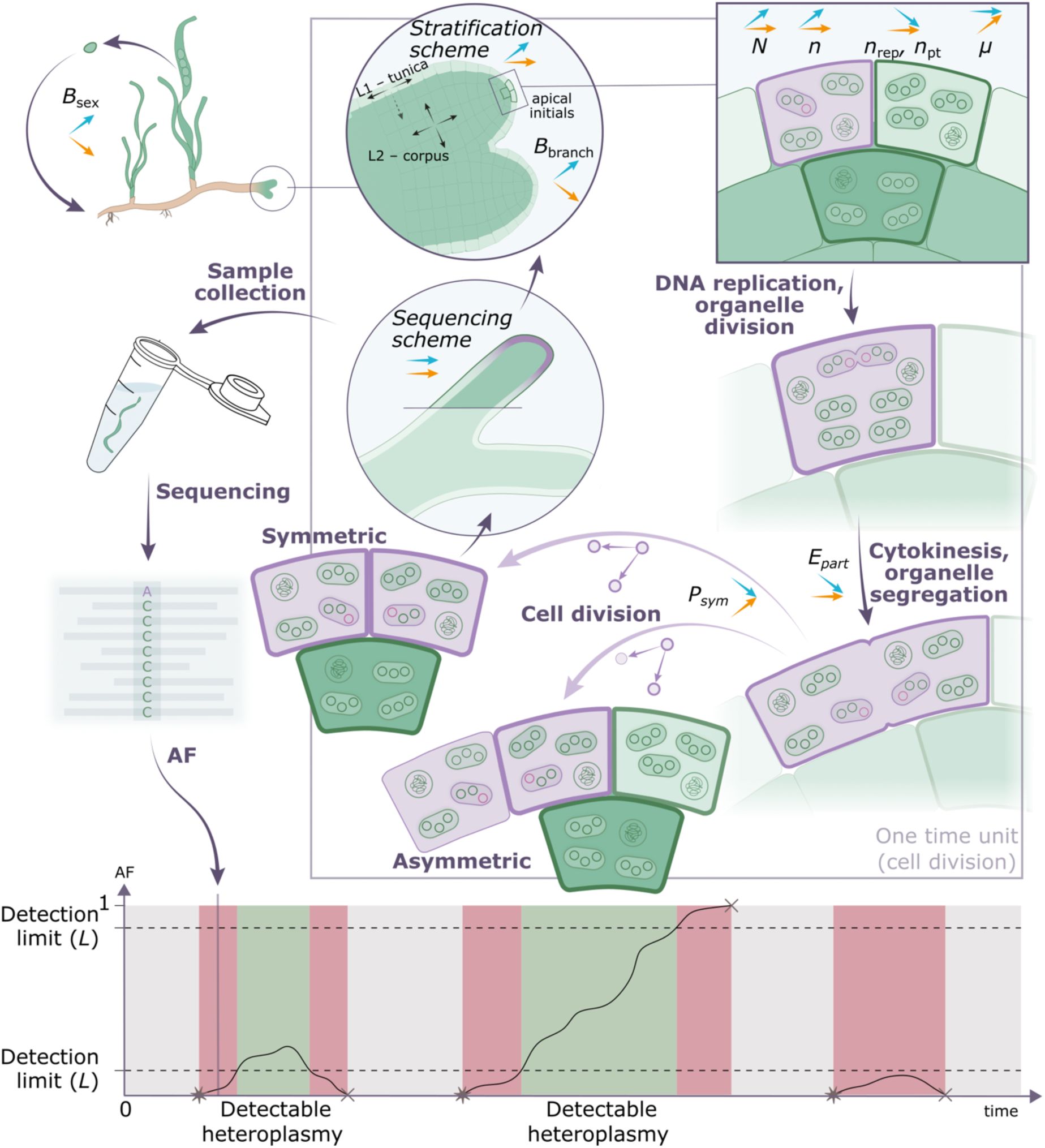
Organelle genome evolution within the stem cells; a model for monocots. The depicted process represents a single stem cell division and highlights the steps and model parameters that affect the result organelle allele composition. The parameters and their descriptions are listed in Table 2. Each parameter is accompanied by two arrows that specify the expected effect of its elevation on substitution rate (orange) and an average neutral allele segregation time (blue). On the bottom graph, the fate of a single locus is followed, where *de novo* mutations arise (stars) and get fixed or extinct (crosses). The model implies that after every circle the tissue is sequenced, revealing the mutant allele frequency in the focal locus that is either detectable (green area on the graph) or undetectable (red area on the graph) given the limitations of the methodology (*L*). The grey area on the graph reflects the homoplasmic state of the system.

As our model species, we chose the seagrass species *Zostera marina* (eelgrass), a widespread marine flowering plant that radiated across the Atlantic and Pacific Oceans several hundred thousand years ago (Yu, Khachaturyan, et al. 2023). Eelgrass reproduces both sexually and vegetatively by branching of rhizomes, which leads to large eelgrass meadow formation by ramets of the same clone that can be several hundred years old (Reusch et al. 1999). The rhizomes involved in ramet initiation are formed from the shoot apical meristem (SAM) (Marbà and Duarte 1998). The availability of the nuclear, plastid, and mitochondrial genome assembly and meristem imaging, together with the uniform worldwide population DNA sequencing dataset, makes *Z. marina* an interesting species for a short-term organelle evolution study (Olsen et al. 2016; Ma et al. 2021; Khachaturyan et al. 2023; Yu, Khachaturyan, et al. 2023; Yu, Renton, et al. 2023). Here, we incorporated the DNA sequencing dataset of *Z. marina* meristematic region samples to study the dynamics of neutral evolutionary processes in mitochondria and plastids (Yu, Khachaturyan, et al. 2023). We examined heteroplasmic loci as the traces of ongoing novel mutation segregation in sequenced samples and used agent-based stochastic computer simulations to estimate the effect of individual factors on the neutral allele emergence and segregation time.

## Results

For the purpose of our study, several definitions are required. Here we use the term ‘population’ to refer to different levels of biological organizations. Primarily, we study the population of organelle DNA molecules (oDNA) – mitochondrial DNA (mtDNA) or plastid DNA (ptDNA) – in a set of SAM initials (Figure 1). Therefore, fixation of a certain allele in the focal oDNA population corresponds to fixation at the level of an individual plant (i.e., substitution), whilst the population of plants might remain heterogeneous. Accordingly, the estimated substitution rate and the number of accumulated mutations take into account all fixation events in one individual plant. We refer to this population as an oDNA, mtDNA, or ptDNA population. Additionally, we addressed the set of shoot apical meristem (SAM) stem cells as the ‘cell population’ when describing processes that manifest at the cellular level. For the conventionally defined populations of individual plants and clones, we used the term ‘eelgrass population’ or, alternatively, specified their geographical location, e.g., ‘Alaskan population’. Thus, we distinguish between effects at different levels of biological organizations with a main focus on the oDNA population genomics.

### Nucleotide substitution rate in the *Z. marina* organelle genomes is homogeneous

To compare the rate of evolution between the two plant organelles, we detected single nucleotide polymorphisms (SNPs) in mitochondrial and plastid genomes across 110 samples collected in nine eelgrass populations residing in the Atlantic Ocean and two populations residing near the Alaskan coast. We focused only on neutral positions (here defined as intergenic positions), in which no SNP detection bias is expected, e.g., by excluding nuclear mitochondrial (NUMTs), nuclear plastid (NUPTs), and mitochondrial plastid (mtptDNA) DNA (Kleine et al. 2009; Scarcelli et al. 2016). The detection of SNPs at neutral positions within the mitochondrial genomes revealed 33-80 SNPs that separate the Atlantic and the Alaskan mitogenomes, further referred to as ‘genetic distance’ (Supplementary Figure S1). The number of SNPs was, on average, 57, which corresponds to an average of 28.5 mutations accumulated since the divergence of the Atlantic and the Alaskan populations, which is estimated to occur 243 kya (Yu, Khachaturyan, et al. 2023). Thus, the total of 28.5 accumulated mutations corresponds to a substitution rate of 9.6 × 10^−10^ substitutions per base pair per year (Table 1). The SNPs at neutral plastid positions revealed a genetic distance of 20-26 SNPs between the Atlantic and the Alaskan plastid genomes (Supplementary Figure S2). The genetic distance was 23 SNPs on average, which corresponds to an average of 11.5 accumulated mutations since the divergence time; 11.5 plastid mutations correspond, thereby, to a substitution rate of 7.9 × 10^−10^ substitutions per base pair per year. That is similar to the estimated mitochondrial substitution rate (Table 1).

**Table 1.** Observed fixed and heteroplasmic SNPs at neutral sites in 163 *Z. marina* samples. Values are means ± standard deviation. Atlantic and Alaskan populations are selected in accordance with the classification of Yu et al. (2023). The total number of heteroplasmic sites per base pair corresponds to the expected heteroplasmy probability (*P*) which is the numeric result of simulation experiments. SSC and LSC abbreviations stay for the small single-copy region and the large-single copy region of the plastid genome correspondingly.

|  | Mitochondrial genome | Plastid genome |
| --- | --- | --- |
| Average genetic distance between the Atlantic and Alaskan populations<sup>1</sup> | 56.7 ± 8.9 | 22.5 ± 1.1 |
| Neutral positions for fixed mutations (per sample) | 121,502 | 58,721 |
| Average number of accumulated mutations since the Atlantic-Alaskan divergence (per base pair)<sup>1</sup> | $2.3 \times 10^{-4} \pm 3.7 \times 10^{-5}$ | $1.9 \times 10^{-4} \pm 9.4 \times 10^{-6}$ |
| Substitution rate (per base pair per year) | $9.6 \times 10^{-10} \pm 1.5 \times 10^{-10}$ | $7.9 \times 10^{-10} \pm 3.9 \times 10^{-11}$ |
| Substitution rate based on the Californian-Pacific divergence (per base pair per year) | $7.1 \times 10^{-10} \pm 8.2 \times 10^{-11}$ | $7.2 \times 10^{-10} \pm 3.7 \times 10^{-11}$ |
| Number of heteroplasmic sites<sup>1</sup> | 1 (+1) | 6 |
| Neutral positions for heteroplasmic sites (total) | 12,920,202 | 3,284,450 |
| Number of heteroplasmic sites per base pair (total)<sup>1</sup> | $7.7 \times 10^{-8}$ <sup>2</sup> | $1.8 \times 10^{-6}$ |
| Heteroplasmic sites<sup>3</sup> | **27884** \| JS03, AF=0.91<br>(+124884 \| FR06, AF=0.51) | SSC:<br>**8108** \| ALI10, AF=0.75,<br>**11673** \| SD08, AF=0.44; SD03, AF=0.48; SD01, AF=0.82;<br>SD07 and SD11 fixed<br>LSC:<br>**83431** \| ALI10, AF=0.78,<br>**98817** \| ALI05, AF=0.50,<br>**118850** \| ALI04, AF=0.94;<br>ALI05, AF=0.50,<br>**120428** \| SD08, AF=0.56 |

<sup>1</sup> At corresponding neutral positions.
<sup>2</sup> For one heteroplasmic site.
<sup>3</sup> Location abbreviations: Japan-South (JS), Alaska-Izembek (ALI), San Diego, California (SD).

An alternative calculation of the substitution rates based on the deepest *Z. marina* divergence between Californian and the Main Pacific populations yields a similar substitution rate for the ptDNA but a markedly lower substitution rate for the mtDNA (Table 1). Unlike the Atlantic and Alaskan populations that are geographically disjoined, gene flow between Californian and Main Pacific populations is likely and was shown for the nuclear genome (Yu, Khachaturyan, et al. 2023). On the one hand, genetic distances based on the California-to-Main Pacific split might underestimate the substitution rate for mtDNA due to effects such as paternal leakage for the *Z. marina* mitogenome, which would be impossible for the plastid genome. On the other hand, however, the clear separation of the Californian haplotypes from the Main Pacific haplotypes points towards a preferential maternal mitochondria inheritance, like for plastids, rather than a biparental inheritance as for the nuclear genome (Supplementary Figures S1, S2; Yu et al. 2023). Therefore, to avoid a possible bias caused by paternal leakage, we based our estimations only on the divergence between the Alaskan and Atlantic populations instead of the deepest divergence between the Californian and main Pacific populations.

Conceptually, the higher copy number of ptDNA compared to mtDNA entails an increase in the mutational supply but a decrease in the probability for neutral allele fixation in a manner that is proportional to the replicon copy number *n* (i.e., the copy number should have an effect akin to population size; reviewed in Lanfear et al. 2014). Therefore, we do not expect a positive association between the replicon copy number and evolutionary rates. Additionally, the allele segregation process itself, when neutral towards mutant alleles, has no effect on the allele fixation probability. In models of single locus dynamics over infinite time (such as the one reported here), genetic drift caused by bottlenecks does not affect the fixation probability of neutral alleles. Nonetheless, mutations arising in the reduced host cell population (just after a bottleneck event) have a higher probability of being fixed. Thus, factors affecting the substitution rate include the mutation rate, the proportion of symmetric cell divisions, and the frequency of bottlenecks. The latter two factors manifest at the cellular level and are, therefore, common to plastids and mitochondria. We conclude that *Z. marina* mitochondrial and plastid genomes have similar substitution rates and, thereby, also similar mutation rates. However, estimating the mutation rate from the substitution rate requires the modelling of the evolutionary process, since multiple factors, such as the proportion of symmetric cell divisions and the frequency of branching and sexual reproduction, affect the fixation probability (Figure 1).

### Establishment of a mathematical model to investigate the determinants of organelle allele dynamics

To further examine the determinants of organelle evolutionary rates, we established a model to simulate neutral allele dynamics in *Z. marina*. The model describes allele dynamics in a single genome locus until a certain number of mutations occur and reach either loss or fixation in the oDNA population. The ultimate goal of the simulation is to calculate the probability (*P*) of detecting heteroplasmy in a focal locus when sequencing plant meristematic region at a random time point (Figure 1). Later, we adjusted the simulation to match it consistent with our empirical findings. The probability *P* is calculated as the proportion of detectable heteroplasmic time span in cell divisions (*T_DET_*) to the total time of the simulation (*T_Total_*). The heteroplasmy was considered detectable if the reference and variant allele frequencies (AFs) were both above a certain frequency threshold *L*, which reflects the sensitivity of the variant detection methodology. Here we assume that the considered neutral genome loci are independent and the *de novo* mutations occur only during the homoplasmic oDNA population state; this assumption is supported by the low mutation rates observed here. Additionally, we assume that the tissue sample is formed of one stem cell descendant and, thus, has the same frequency of the mutant allele as the stem cell itself regardless of the mutant copy distribution among cells (similar to recently published simulations by Broz et al. 2023). We also assume no *de novo* mutations during tissue formation since the probability for such mutations to arise and reach the detectable AF is low. Thus, the two major components that contribute to the probability *P* are the mutational supply and the average detectable allele segregation time *T_DET_*.

In our model, we consider a set of meristem stem cells to have a constant size of 20 cells organized as a cell bulk with two layers – the tunica layer (L1) and the corpus layer (L2) – each consisting of two rows of five cells. The oDNA copy numbers in stem cells were set according to our estimates of the average mtDNA and ptDNA abundance across 24 samples of the *Z. marina* meristematic region (Supplementary Table S1). Based on those estimates, we assume that each stem cell contains 40 copies of mtDNA and 216 copies of ptDNA (Table 2). Note that previous studies point towards a slightly lower ptDNA copy number, e.g., 35-192 copies per cell in beet, *A. thaliana*, tobacco, and maize, and slightly lower mtDNA copy number, e.g., up to 65 copies per cell in maize (Kumar et al. 2014; Greiner et al. 2020). Time units in the model correspond to the duration of one cell division in the population of stem cells (Figure 1). Cell divisions are accompanied by several stochastic processes: (i) the random sampling of the next cell to divide (with equal probabilities for all stem cells), (ii) the choice between symmetric and asymmetric cell division (model parameter *P_sym_*), (iii) the random sampling of the cell to be removed from the cell population in case of a symmetric cell division (the probabilities are defined by the model parameter *stratification scheme*), and (iv) the replicated organelle allele segregation to the two daughter cells (that can be random, or defined by the parameter *E_part_*). Additionally, we applied regular bottlenecks of one corpus cell to mimic sexual reproduction events and of ten cells – two vertically stacked rows one from tunica and one from corpus – to mimic branching events. The frequency of the two bottleneck types is defined by the model parameters *B_sex_* and *B_branch_*, respectively (Table 2). Note that the frequency of branching bottlenecks (*B_branch_*) might differ from the frequency of branching events as only one of the two shoot axes undergoes subsampling of stem cells when the bifurcation is lateral. For dichotomous bifurcation, each branching event leads to a cell bottleneck on both axes (reviewed in Müller and Leyser 2011; Gola 2014). After a bottleneck, the cell population is assumed to proliferate until the given constant cell population size is reached.

**Table 2.**
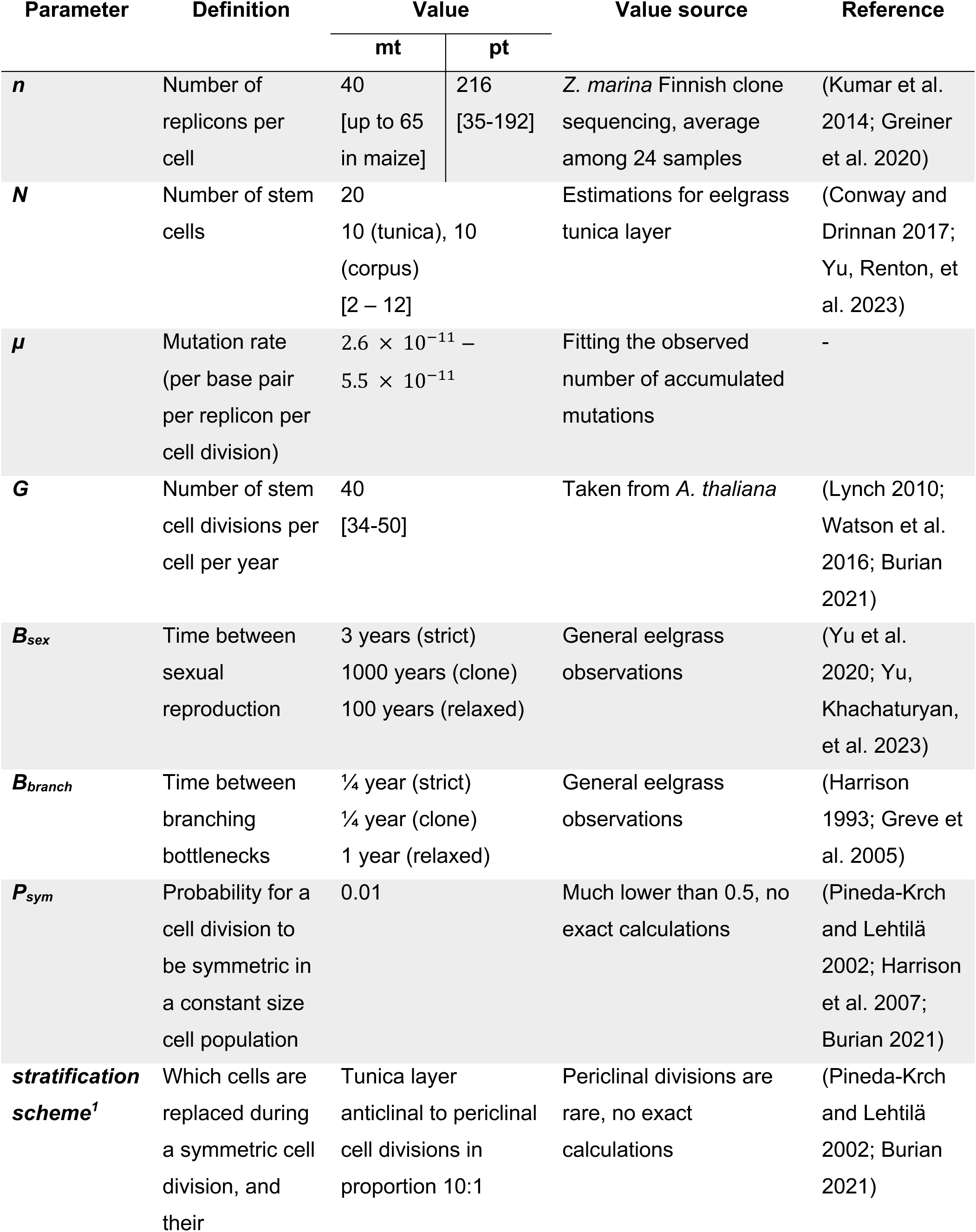
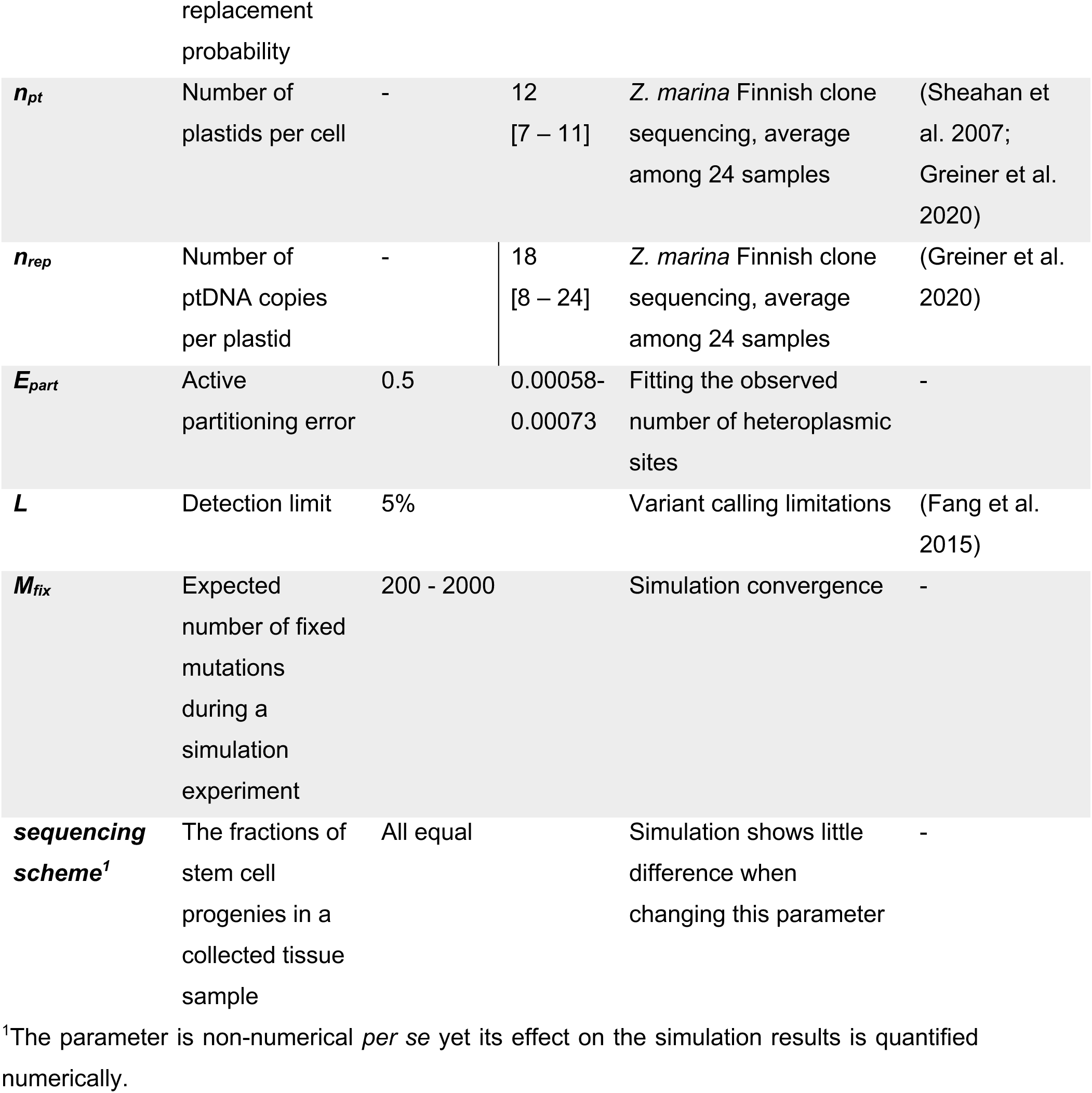
Parameters of the agent-based model. The values used in the simulations for mitochondria (mt) and plastids (pt) were taken from literature and *Z. marina* experiments when possible. Numbers in square brackets show the ranges based on existing data.

| Parameter | Definition | Value |  | Value source | Reference |
| --- | --- | --- | --- | --- | --- |
|  |  | mt | pt |  |  |
| ***n*** | Number of replicons per cell | 40<br>[up to 65 in maize] | 216<br>[35-192] | *Z. marina* Finnish clone sequencing, average among 24 samples | (Kumar et al. 2014; Greiner et al. 2020) |
| ***N*** | Number of stem cells | 20<br>10 (tunica), 10 (corpus)<br>[2 – 12] |  | Estimations for eelgrass tunica layer | (Conway and Drinnan 2017; Yu, Renton, et al. 2023) |
| ***μ*** | Mutation rate (per base pair per replicon per cell division) | $2.6 \times 10^{-11}$ –<br>$5.5 \times 10^{-11}$ | | Fitting the observed number of accumulated mutations | - |
| ***G*** | Number of stem cell divisions per cell per year | 40<br>[34-50] |  | Taken from *A. thaliana* | (Lynch 2010; Watson et al. 2016; Burian 2021) |
| ***B<sub>sex</sub>*** | Time between sexual reproduction | 3 years (strict)<br>1000 years (clone)<br>100 years (relaxed) |  | General eelgrass observations | (Yu et al. 2020; Yu, Khachatryan, et al. 2023) |
| ***B<sub>branch</sub>*** | Time between branching bottlenecks | ¼ year (strict)<br>¼ year (clone)<br>1 year (relaxed) |  | General eelgrass observations | (Harrison 1993; Greve et al. 2005) |
| ***P<sub>sym</sub>*** | Probability for a cell division to be symmetric in a constant size cell population | 0.01 |  | Much lower than 0.5, no exact calculations | (Pineda-Krch and Lehtilä 2002; Harrison et al. 2007; Burian 2021) |
| ***stratification scheme<sup>1</sup>*** | Which cells are replaced during a symmetric cell division, and their replacement probability | Tunica layer<br>anticlinal to periclinal cell divisions in proportion 10:1 |  | Periclinal divisions are rare, no exact calculations | (Pineda-Krch and Lehtilä 2002; Burian 2021) |
| $n_{pt}$ | Number of plastids per cell | - | 12<br>[7 – 11] | *Z. marina* Finnish clone sequencing, average among 24 samples | (Sheahan et al. 2007; Greiner et al. 2020) |
| $n_{rep}$ | Number of ptDNA copies per plastid | - | 18<br>[8 – 24] | *Z. marina* Finnish clone sequencing, average among 24 samples | (Greiner et al. 2020) |
| $E_{part}$ | Active partitioning error | 0.5 | 0.00058-0.00073 | Fitting the observed number of heteroplasmic sites | - |
| $L$ | Detection limit | 5% | | Variant calling limitations | (Fang et al. 2015) |
| $M_{fix}$ | Expected number of fixed mutations during a simulation experiment | 200 - 2000 | | Simulation convergence | - |
| **sequencing scheme<sup>1</sup>** | The fractions of stem cell progenies in a collected tissue sample | All equal |  | Simulation shows little difference when changing this parameter | - |

<sup>1</sup>The parameter is non-numerical *per se* yet its effect on the simulation results is quantified numerically.

### The mutation rate ranges between ca. 2. 6 × 10^−11^ – 5. 5 × 10^−11^ mutations per base pair per replicon per cell division in both organelles

In order to estimate the oDNA mutation rates (*μ*), we simulated the allele dynamics for different *μ* values to calculate the expected number of substitutions in a single locus in 243,300 years. This result, multiplied by the number of sites, thereby corresponds to the expected number of accumulated mutations in a *Z. marina* individual since the Alaskan-Atlantic divergence. Under a neutral regime, the mutation rate equals the substitution rate (Kimura 1968). Indeed, we found the fixation rate and the mutation rate *μ* to be linearly dependent, enabling us to estimate the mutation rate, *μ*, from the value that best fits the observed number of fixed mitochondrial and plastid mutations (Figure 2A). Bottleneck regimes with high branching rates, ‘strict’ and ‘clone’, and low branch rates, ‘clone’ and ‘no’, form separate clusters, hence the frequency of branching had a major effect on the estimated *μ* when applying different bottleneck regimes. However, the total range of the estimated mutation rate remained within a narrow range of 4.3 × 10^−11^ – 4.7 × 10^−11^ and 3.5 × 10^−11^ – 3.9 × 10^−11^ mutations per base pair per replicon per cell division for mtDNA and ptDNA, respectively (Figure 2A). Likewise, the proportion of symmetric cell divisions (*P_sym_*) had a minimal effect on the estimated mutation rate, *μ*, if considered within a realistic range 0.1-0.005 (Supplementary Figure S3). To transform the unit of time (total number of cell divisions) used in our model into time elapsed in years, we used a constant cell depth, *G* = 40 cell divisions per stem cell per year, as for an annual plant *A. thaliana*. Although eelgrass might refrain from sexual reproduction for tens of years, eelgrass ramets, unlike tree branches, separate from their parental plants and therefore are susceptible to selection within a single clone. Hence, we assume that the number of stem cell divisions per year in eelgrass is similar to those of annual plants. The mutation rate is proportional to the cell depth, *G*, which is unlikely to assume a value outside the range of 34-50 cell divisions per stem cell per year (reviewed in Burian 2021). Taken together, we infer that the absolute mutation rate of *Z. marina* mitochondria is between 3.2 × 10^−11^ and 5.5 × 10^−11^ and in plastids it is between 2.6 × 10^−11^ and 4.7 × 10^−11^ mutations per base pair per replicon per cell division (note that a rough estimate of the mutation rate can be achieved also by dividing the substitution rate by *G*).

**Fig. 2:**
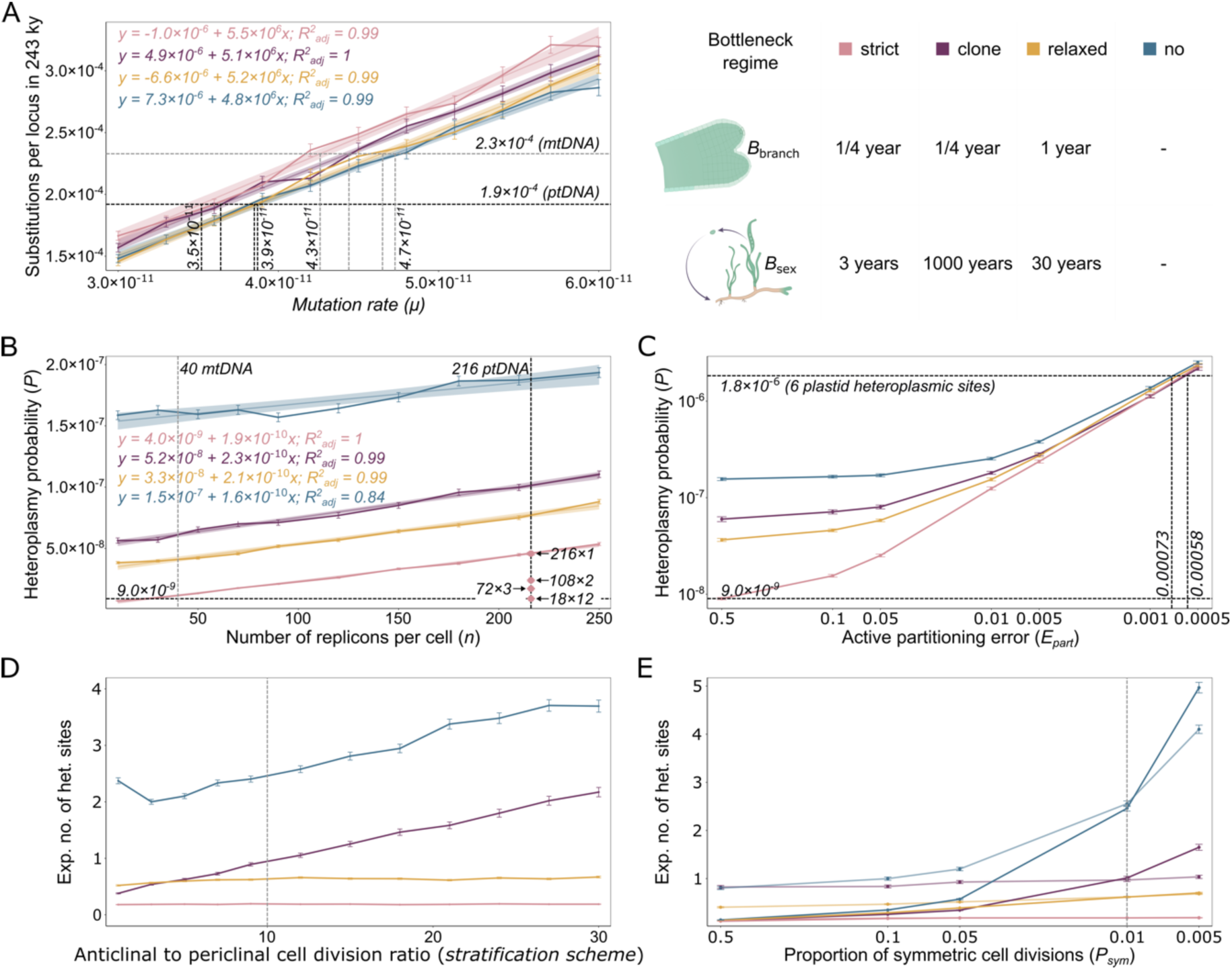
The effect of individual parameters in the simulation of *Z. marina* evolution of organelle genomes in the absence of selection. a, The number of mitochondrial substitutions per locus in 243,300 years in the *in-silico* experiments for different mutation rates (*µ*). The dashed grey (mitochondrial genome) and black (plastid genome) lines indicate the *µ* values matching the observed number of accumulated fixed mutations. The legend shows the colors and the parameters used for the four bottleneck regimes in Figure 2,3. b, The alteration of the heteroplasmy probability (*P*) with an increase in the replicon copy number (*n*) for *µ* that corresponds to the plastid genome. Points show the estimated *P* when applying the plastid hierarchical structure in the range of 216 ptDNA/pt for 1 pt/cell to 18 ptDNA/pt for 12 pt/cell. Horizontal dashed lines indicate the *P* values for mtDNA (grey for mitochondrial *µ*, black for plastid *µ*) and ptDNA (black) with the strongest hierarchy. c, The estimated probability *P* when an active plastid segregation is applied with a certain active partitioning error (*E_part_*). Horizontal dashed grey lines indicate the *P* value for a random partitioning and the expected *P* based on the observation data. d, The dependency of the expected number of mitochondrial heteroplasmic sites in the worldwide dataset when increasing the tunica and corpus layer isolation. The vertical dashed line indicates the default stratification parameter. e, The effect of the symmetric cell division proportion (*P_sym_*) with the fixed level of stratification (bold lines) and adjusted stratification level (transparent lines) on the expected number of mitochondrial heteroplasmic sites. The vertical dashed line indicates the default *P_sym_* parameter.

### Allele segregation is ca. 20-fold slower in plastids compared to mitochondria

Comparing the segregation times of neutral alleles between mitochondria and plastid genomes requires a comparison of the number of heteroplasmic loci that may reveal ongoing allele segregation across multiple sequenced samples. For that purpose, we identified all heteroplasmic SNPs at neutral positions for heteroplasmic sites across 163 *Z. marina* samples collected in 16 locations of the Pacific and Atlantic Oceans (Table 1). In total, one heteroplasmic locus was detected in mitochondrial genomes with one additional heteroplasmy of an unclear origin (Supplementary Note S1). Six heteroplasmic loci were detected in the plastid genomes of the 163 samples. Since the total number of neutral positions for heteroplasmic sites is 3.9 times higher for mitochondria compared to plastids (Table 1), we conclude that heteroplasmy is 23 times more abundant in the plastid compared to the mitochondrial genome.

As a next step, we used our model to identify the factors that would be able to explain such a dramatic difference in observed heteroplasmy. We focused on three factors that distinguish mitochondrial and plastid allele segregation dynamics: replicon copy number, hierarchical structure, and the mode of segregation (i.e., partitioning mechanism). The simulation demonstrates that when the replicon copy number increases from 40 (*n_mtDNA_*) to 216 (*n_ptDNA_*), *P* increases 1.2-3.9 times depending on the bottleneck regime (Figure 2B), that is much lower than the observed 23-fold difference in the number of heteroplasmic sites. Notably, the simulation results show a linear association between heteroplasmy probability and the copy number. The linear association is well explained by the constant total number of cell divisions in a single simulation experiment and the average time of detectable heteroplasmy (*T_DET_*) that is nearly constant in the simulation, such that the increase in copy number affects solely the mutational supply, resulting in a linear dependency (Supplementary Figure S4AB). The observation of nearly constant detectable segregation time implies that the increased segregation time is being buffered by the increased minimum number of mutant replicons that pass the detection limit threshold *L*. Furthermore, the hierarchical plastid structure significantly decreases the segregation time, which results in a heteroplasmy probability for ptDNA that is even slightly lower than for mtDNA (Figure 2B, see point for 18x12, Supplementary Figure S4C). Thus, the third factor – the mode of segregation – is the only one remaining to explain the difference in the number of observed heteroplasmic sites. Here we assume that nuclear control of equal plastid segregation to different daughter cells in stem cells occurs via an active partition of sister plastids after plastid division. In the extreme scenario, such a mode of segregation never leads to allele fixation in the absence of biparental inheritance, as shown in the example of nuclear mutations in clones (Yu et al. 2020). However, the fixation of novel alleles is possible if we allow a certain probability of partitioning error. A partitioning error of 0.5 indicates that the two sister plastids segregate to the same daughter cell with a probability of 0.5, thus simulating random segregation. The simulation demonstrates that a partitioning error of 0.00058-0.00073 results in a heteroplasmy probability *P* that is consistent with the observed number of ptDNA heteroplasmic sites in the sequencing data (Figure. 2C). Thus, we conclude that the observed difference between plastid and mitochondrial allele dynamics is likely explained by a relatively strict active partitioning of plastids during stem cell division, very much in contrast to the random segregation of mitochondria. Notably, when applying strict active partitioning, plastids become homoplasmic considerably fast (within 740-1140 cell divisions, on average, depending on the bottleneck regime, which corresponds to 0.9-1.4 years), while the heteroplasmy is maintained in the oDNA population on the level of plastids in cells, as cells reach homoplasmy 300-550 times longer. Consequently, the number of plastids per cell has a minimal effect on the segregation time, if the number of ptDNA copies per plastid remains constant (Supplementary Figure S4E). In contrast, when applying random plastid segregation (*E_part_*, = 0.5), the average time until the first homoplasmic plastid occurs (560-740 cell divisions, which corresponds to 0.7-0.9 years) is similar to the time until the first homoplasmic cell occurs (1200-1700 cell divisions, which corresponds to 1.5-2.4 years).

### Meristem stratification increases allele segregation time in the absence of frequent sexual reproduction

In the next step, we performed simulations to evaluate the effect of four parameters that are acting on the cellular level and, therefore, are common for both organelles: *stratification scheme*, *P_sym_*, *sequencing scheme*, and the number of stem cells (*N*). To evaluate the effect of the *stratification scheme* in the model we applied different ratios of the anticlinal to periclinal cell division in the tunica layer, which reflects stratification between the tunica (L1) and the corpus (L2) meristematic layers. Periclinal divisions were assumed to occur only during the non-proliferating stage and only in the direction from tunica to corpus. The simulation results predict a strong impact of the frequency of sexual reproduction on the effect of stratification since the regular single-cell bottlenecks inevitably admix the genetic content of the two meristematic layers such that the effect of stratification is limited (Figure 2D, bottleneck regimes ’strict’ and ’relaxed’). For the bottleneck regimes featuring the clonal lifestyle, however, a relatively strict genetic isolation of the two layers leads to an increase in the expected number of heteroplasmic sites (Figure 2D, bottleneck regimes ’clone’ and ’no’). Decreasing the parameter *P_sym_*, proportion of symmetric cell divisions, significantly increases the probability *P* and thereby the expected number of heteroplasmic sites (Figure 2E). However, the major contribution to the observed elevation arises from the corresponding decrease of the periclinal genetic material flow from the tunica to the corpus. The individual effect of the parameter *P_sym_* is minimal and is buffered by branching bottlenecks that inevitably imply that cells are replaced during the proliferation stage (see the effect of stratification in Figure 2E). Furthermore, we considered one factor of the system that does not affect the segregation dynamics, but might alternate the estimated AF – the proportion of the stem cell progenies in the sequenced tissue, *sequencing scheme*. The sequenced meristematic region contained early-developed tissues, such as leaf primordia. Since the periclinal division is common in leaf formation, the proportion of tunica and corpus progenies is difficult to estimate. We conducted the calculations assuming the proportion of the tunica layer progenies to the corpus layer progenies from 16 to 1/16. From the low variance of the estimated expected number of heteroplasmy sites for all bottleneck regimes that had any bottlenecks, i.e., except for ‘no’, we concluded that disproportional contribution of the stem cells to the sequenced tissue has a minimal effect (Supplementary Figure S4D). Therefore, by default we used an equal number of progenies for all stem cells in our simulations by default. Similarly, the number of stem cells, *N*, has a minimal effect on both the allele fixation and the allele segregation processes unless no bottlenecks are applied (Supplementary Figure S5).

If we use an extreme combination of the simulation parameters *stratification scheme* of 30:1 (anticlinal to periclinal cell divisions) and *P_sym_* = 0.005 and ignore bottlenecks, the expected number of heteroplasmic sites in 163 samples is 7.8 for mitochondria and 1.7 for plastids. Notably, under these extreme parameters, the estimated number of heteroplasmic sites for the mitochondria is considerably higher than the observed one (or two) sites in the sequencing data, whilst the estimation for plastids remains significantly lower than the observed six heteroplasmic sites. Note that we considered stochastic partitioning, *E_part_* = 0.5. Hence, our analyses of the effects of parameters support our hypothesis that the partitioning of sister plasmids to the daughter cell is not stochastic.

### Plastids but not mitochondria are expected to have common heteroplasmic sites between different clones of the same eelgrass population

To evaluate the limitation of the whole genome sequencing approach for detecting heteroplasmy, we applied different detection limits *L*. Note that the total number of ptDNA copies in the focal oDNA population in the simulation is 4,320 (216 *copies per cell* × 20 *cells*) and the total number of mtDNA is 800 (40 *copies per cell* × 20 *cells*). A threshold of zero features a noiseless allele detection process with one mutant copy to be sufficient for heteroplasmy detection, whilst the highest threshold of 0.2 requires at least 864 plastid and 160 mitochondrial mutant copies for the detection. Our results demonstrate that a 10-fold decrease in the detection limit from the default value of 0.05 to 0.005 only shows a slight increase in the expected number of heteroplasmic sites for both mitochondria and plastids (Figure 3A). Yet, the spike for *L*=0 shows the maximum number of heteroplasmic sites in 163 samples to be 5.6 for mtDNA and 14.2 ptDNA. Thus, the number of novel mutations that will eventually become fixed (or are lost) in a single sampled eelgrass individual is expected to be below one. Consequently, we conclude that if an organelle heteroplasmy is detected in a single sequenced individual, it is unlikely to refer to an ongoing segregation of a novel mutation.

**Figure 3:**
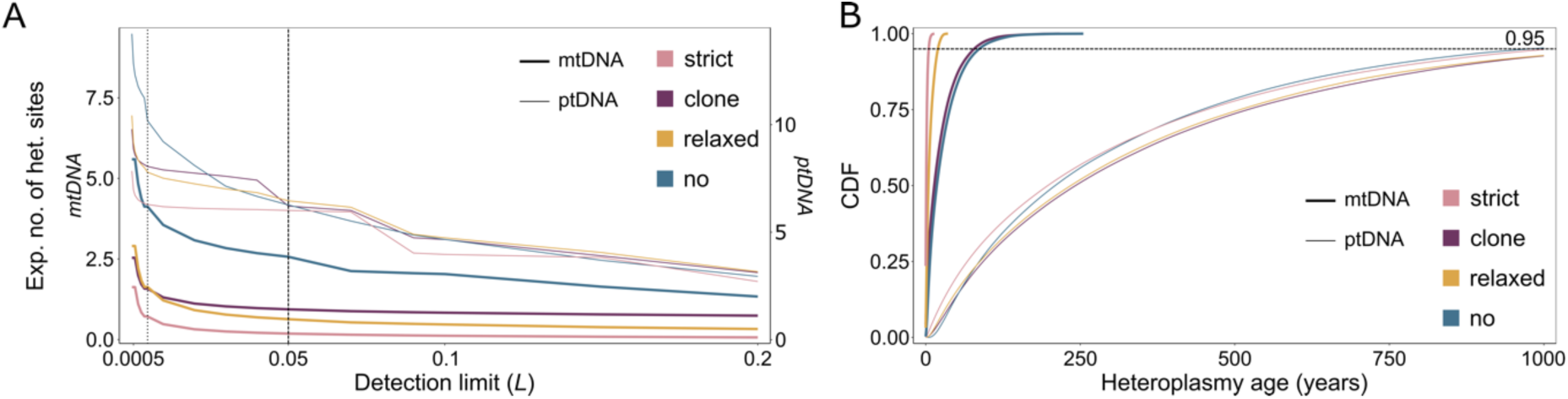
Allele dynamics in a single simulation experiment of *Z. marina* mtDNA and ptDNA neutral evolution. a, The expected number of heteroplasmic sites in 163 samples in mitochondrial (bold lines) and plastid (thin lines) genomes for different heteroplasmy allele frequency detection limits (*L*). The vertical dashed line indicates the default detection limit of 0.05, the vertical dotted line indicates the detection limit of 0.005. b, The cumulative age distribution of detectable heteroplasmy in mitochondria (bold lines) and plastids (thin lines). Colors correspond to the bottleneck regimes in accordance with Figure 2.

To this end, we used the selected simulation parameters that are in accordance with plant biology and the observed data to estimate the age distribution of detectable heteroplasmic sites in mitochondria and plastids. The simulation indicates that while 67-73% of the detectable heteroplasmic sites in plastids are over 100 years old and 5-7% are over 1000 years old, only 2-3% of mitochondrial heteroplasmic sites persist for at least 100 years given a clonal lifestyle (Figure 3B). Eelgrass populations with regular sexual reproduction are not expected to maintain intra-individual mitochondrial heteroplasmy for over 36 years (Figure 3B). Our prediction that old heteroplasmy can be found in the plastid genome is confirmed by the empirical data. Two out of six detected heteroplasmic sites in the plastid genome – position 11673 in the San Diego population and position 118850 in the Alaska-Izembek population – are heteroplasmic at the level of individual plants. At the same time, these two sites have a fixed mutant genotype in five (position 11673) and two (position 118850) samples taken from the same location (Table 1). Given that all the samples belong to different clones, the mutations likely emerged in the ancestor of the sampled clones, thereby persisting in the eelgrass population for a significant time and surviving at least one single-cell bottleneck. The presence of heteroplasmy in multiple clones in the plastid genome but not in the mitochondrial genome provides an additional indirect confirmation for our conclusion that the segregation time of plastid alleles is significantly elongated.

## Discussion

Studying the evolution of organelles is crucial for our understanding of plant adaptation to changing environments due to their contribution to cellular respiration, photosynthesis, and a myriad of other essential processes. Here, we show how the organization and partition of the plant organelles within the germline affect the evolution of their genomes. In doing so, we uncover the fundamental principles of organelle population genetics that are essential for further investigations of long-term evolution and molecular dating of divergence events.

Mutation rates are generally estimated for nuclear genomes and in units per plant generation (Krasovec et al. 2018; Zhang et al. 2018). More rarely, the mutation rate is given per stem cell division, which allows comparison across species; this rate might be as low as 1 × 10^−11^ in lotus (*Nelumbo Adans.)* in comparison to 1.6 × 10^−10^ in *Arabidopsis thaliana* (Lynch 2010; Ossowski et al. 2010; Zheng et al. 2022). Since organelle genomes are much smaller than the nuclear genome (usually < 400 kb), estimates of evolutionary rates based on mutation accumulation require observations over a long-time scale. The majority of estimations for the plant organelle evolutionary rate are based on synonymous and nonsynonymous substitutions in cross-species comparisons (Wolfe et al. 1987; Muse 2000; Drouin et al. 2008; Xu et al. 2012). These studies consistently show a significantly faster plastid gene evolution in comparison to mitochondrial gene evolution (Wolfe et al. 1987; Drouin et al. 2008). Individual analysis of plastid genomes showed inverted repeats to evolve slower compared to single copy loci, similar to mitochondria (Muse 2000). However, only a few studies measured the evolutionary rates of mitochondrial and plastid genomes for individual species. One example is a previous study of Douglas fir (*Pseudotsuga menziesii*) that shows a similarity between the two organelle substitution rates of 5.26 × 10^−10^ and 4.41 × 10^−10^ substitutions per base pair per year for mtDNA and ptDNA, respectively (Gugger et al. 2010), which are slightly lower compared to eelgrass (Table 1). The similarity between mitochondrial and plastid substitution rates is concordant with the similarity in their core enzymes and components of the oDNA replication system, which are identical for plastids and mitochondria in land plants (Moriyama and Sato 2014). Why studies based on interspecies comparison result in significantly different mutation rates between the two land-plant organelles, while the single-species studies demonstrate the similarity of mutation rates, remains a conundrum. One possible explanation is a long-term effect of DNA recombination, which may be strong for mtDNA and inverted repeats in ptDNA, but not as strong for ptDNA single-copy loci (reviewed in Maréchal and Brisson 2010). Notably, all observed plastid heteroplasmic sites were located in single-copy regions (Table 1). Hence, the neutral allele segregation time in the inverted repeats is likely to be shorter than in the single copy regions. The observed difference might be a direct consequence of the frequent recombination of the inverted repeats, which increases genetic drift and thereby accelerates the segregation of plastid alleles.

Both the observed number and the model prediction of heteroplasmic sites are much lower than the number of samples even if all variants are detected, i.e., the detection limit has no effect. This suggests that detecting any heteroplasmy in a single sequencing experiment is unlikely. To find sufficient heteroplasmic sites in organelles one would need to sequence multiple samples from multiple populations. This conclusion agrees with Scarcelli et al. (2016), who suggested that all plastid heteroplasmic sites in sequencing data are likely to refer to shared DNA fragments among plastids, mitochondria, and the nucleus (i.e., NUMTs, NUPTs, and mtptDNA) that quickly accumulate mutations after transfer (Michalovova et al. 2013; Sloan and Wu 2014). Although we excluded shared DNA regions from the analysis of neutral positions for heteroplasmic sites, the most abundant false positive heteroplasmy pattern includes positions that are clustered together and commonly have a high read coverage. Since the variant allele frequency in these loci was associated with high coverage, they are considered to have a low level of confidence. An alternative explanation is recently integrated NUMTs and NUPTs. In addition to intra-individual polymorphisms created by shared DNA, we observed that mitochondrial small imperfect repeats may generate false positive heteroplasmy signals. At the same time, considering one additional mtDNA heteroplasmic position to be genuine (Table 1) would not change the main conclusion we draw from the observation data. The difference between the plastid and the mitochondrial heteroplasmy abundance thereby would be 12-fold, which we explain by the difference in the partitioning mechanism. Moreover, since the plastid heteroplasmy is expected to be observed in closely related clones, our assumption that the sequenced samples are independent may lead to an overestimation of the expected number of heteroplasmic sites from the heteroplasmy probability *P*. The expected number of heteroplasmic sites is likely to be overestimated in mitochondria as well due to regular mtDNA recombination that was not considered in our simulations. Notably, a study of different date palm cultivars shows a similar pattern; all detected mitochondrial heteroplasmic sites are unique for the corresponding cultivars, while all of the plastid sites are heteroplasmic in all of the cultivars (Sabir et al. 2014). This observation suggests that the dramatic difference in the allele segregation time for mitochondria and plastids might not be a unique characteristic of eelgrass but rather a common evolutionary principle across plant taxa. Our simulation proposes an active partitioning of plastids during cell division. To our knowledge, the fate of sister plastids following stem cell division has not been shown experimentally in any plant species. However, evidence has been presented for the participation of actin filaments and for the co-dependence of the cell and plastid cycles, suggesting a non-stochastic partitioning (Sheahan et al. 2004; Chen et al. 2009; reviewed in Pedroza-Garcia et al. 2016). The nuclear control over plastid division is stronger in meristematic cells (Swid et al. 2018). In monoplastidic organisms, such as basal-branching algae, the plastid and nuclei division and the partitioning during the cytokinesis are well coordinated (Sumiya et al. 2016). Previously, it has been suggested that the evolution of a nuclear control of the plastid cycle in land plants enabled the switch from monoplastidity to polyplastidity (de Vries and Gould 2018). Yet, the mechanisms of plastid copy number control in stem cells are likely to remain unchanged. Consequently, the relocation of plastids from the perinuclear area to the daughter newly formed nuclei is likely to be achieved by active pulling of sister plastids to the opposite poles. Although active partitioning is understudied for eukaryotic extrachromosomal genetic elements, mechanisms of active partitioning are widely described for plasmid segregation in bacteria (reviewed in Bouet et al. 2007; Baxter and Funnell 2015).

In our simulations, we estimated the individual effects of multiple parameters on the dynamics of allele segregation. The fixation time in mtDNA populations depends primarily on the rate of the mutation propagation to other cells and meristematic layers. Therefore, the stratification significantly extends the mitochondrial allele segregation time given that the population undergoes few single-cell bottlenecks (i.e., with rare sexual reproduction events). This result is in agreement with studies showing that stratification and cell-to-cell competition decrease the accumulation of deleterious mutations (Pineda-Krch and Lehtilä 2002; Klekowski 2003; Edwards et al. 2021). A higher number of stem cells does not result in an increased segregation time. Thus, the major drivers of the mutation propagation within a meristematic layer are likely bottleneck events during stem branching and seeding, which are followed by cell proliferation until the total number of stem cells is reconstituted. In contrast, if we indeed assume, as the model suggests, the presence of an active partitioning mechanism for plastids, the limiting process in the new plastid allele segregation is the within-cell fixation rather than the mutation propagation among cells. In that case, the key parameter leading to the extension of plastid allele segregation time is the plastid partitioning error (*E_part_*). Our model shows that the fixation of new mitochondrial variants is achieved relatively fast at all levels of organization, including individual organelles and cells, separate branches, and individual plants. The presence of mitochondrial genetic diversity at multiple levels implies that purifying selection may act at different organizational levels as well. In contrast, plastid variants are fixed within individual organelles, yet the plastid genome remains heteroplasmic at all higher levels of organization over a long period of time, and that may hinder purifying selection against slightly deleterious plastid alleles. Unlike mitochondria, whose function is relatively uniform across tissues, proplastids in meristematic cells perform only basal metabolic functions. We speculate that potentially deleterious or beneficial plastid mutations may be neutral at the level of meristematic cells. Consequently, the competitive pressure among stem cells might be weaker for slightly deleterious plastid variants than for mitochondrial variants, even if allele fixation is reached at the cellular level. How the presence of organelle heteroplasmy at different levels of organization figures into plant adaptation remains a matter for further investigation.

Notably, the effect of drift and selection at multiple levels of organization has been recognized in other extrachromosomal genetic elements (ECEs), e.g., prokaryotic plasmids (Ilhan et al. 2019). Indeed, efforts to model ECE dynamics and evolution typically encounter similar challenges (e.g., Hunter and Fusco 2022; Santer et al. 2022; Broz et al. 2023). Nonetheless, despite common evolutionary characteristics, research on the biology and evolution of oDNA and other ECEs is typically disjoined due to disciplinary boundaries. The development of evolutionary theories and mathematical models of ECE population genetics supplies a basis for cross-disciplinary evolutionary research.

## Materials & Methods

### Data source

Chromosome-level nuclear and plastid *Z. marina* genome assembly were downloaded through the ORCAE platform (https://bioinformatics.psb.ugent.be/gdb/zostera/; Ma et al. 2021). The complete mitochondrial genome assembled in Khachaturyan et al. (2023) was obtained from the NCBI (accessions: OR336317-OR336318). The Illumina whole-genome sequencing of *Z. marina* meristematic region of the inner leaf base was obtained from JGI, Proposal ID: 503251, for the worldwide population dataset and from the NCBI (BioProject: PRJNA557092) for the Finnish clone at Ängsö dataset (Yu et al. 2020; Yu, Khachaturyan, et al. 2023). Worldwide population dataset samples were preselected according to Yu et al. (2023) to avoid representatives of the same clone and an effect selfing on detected variants.

### Detection of genetic variants

We used the *Zostera marina* whole-genome sequencing dataset that comprises 163 samples of meristematic region tissue collected from individual clones in 16 locations of the Atlantic and Pacific Oceans (Yu, Khachaturyan, et al. 2023). Raw Illumina short reads of the 163 samples were aligned to the reference plastid and mitochondrial genomes by BWA-MEM with default parameters and processed by SAMtools v1.10. Allele frequencies were calculated by pysamstats (Li 2013; Danecek et al. 2021; https://github.com/alimanfoo/pysamstats). Additionally, variants were called by BCFtools mpileup v1.8 with unlimited alignment depth and the --count-orphans parameter (Danecek et al. 2021). DNA regions shared between mitochondria and nucleus (NUMTs), or plastids and nucleus (NUPTs), or mitochondria and plastids (mtptDNA) were identified with BLAST (Altschul et al. 1990) of the reference genome pairs with parameters previously used by Smith et al. (2011). In order to avoid undetected shared DNA regions and misaligned repeats, a proportion of the nucleotide position alignment depth from the median coverage of the corresponding genome segment was calculated for each position in the mitochondrial genome. Genome segments were annotated in accordance with the mitogenome rearrangements. Detected NUMTs and mtptDNA were excluded from the calculations (see also Khachaturyan et al. 2023). The coverage proportion of positions in plastid genomes was normalized to either the single copy or inverted repeat area.

For the classification of neutral positions in the mitochondrial and plastid genomes where fixed mutations or heteroplasmic sites were detected, we applied the following filtration steps: (i) DNA regions shared among mtDNA, ptDNA, and the nuclear genome were excluded from the calling of fixed or heteroplasmic variants due to the possibility of identifying potential mutations on other replicons; (ii) microsatellite areas were excluded as these are recognized hotspots for read misalignment; (iii) positions that were significantly under- or overcovered by sequencing reads in comparison to the average coverage of the corresponding genome region were excluded. For the identification of the fixed mutations and heteroplasmic sites, we additionally applied thresholds of allele frequency (AF) and relative coverage, followed by a manual inspection of all remaining variants (see Supplementary Note S1 for detailed information). The genetic distance between two location groups (Alaskan and Atlantic, Californian and main Pacific) was calculated as the average genetic distance in neutral fixed single nucleotide polymorphisms (SNPs) between each pair of samples collected in the corresponding locations.

### Simulation

Stochastic agent-based computer simulations that implement the mitochondrial and plastid neutral evolution model were performed with code in Python v3.10.5, the scrips are available at https://github.com/zyukeriya/plantOrganelleEvolSim. For each parameter set, the simulation was run until the number of mutations that arose in the oDNA population reached a threshold of *n* × *N* × *M_fix_*, where *n* is the replicon copy number per cell and *N* is the number of stem cells. *M_fix_* is a constant that was arbitrarily set as 2,000 for the mutation rate calculation experiments, 500 for the mitochondrial evolution simulations, and 200 for the plastid evolution simulations. The time between mutation events was drawn from a geometric distribution with the success (mutation) probability:

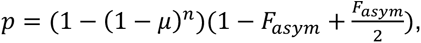

where the fraction of asymmetric cell divisions is:

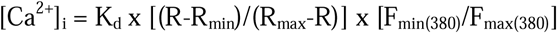

*μ*, *n*, *B_sex_*, *B_branch_*, *N*, and *P_sym_* are defined in Table 2. The estimation error was calculated as the standard error of the mean for the distribution formed by the outcomes of individual mutations. Effects of the *sequencing scheme*, detection limit (*L*), and the distribution of heteroplasmy age were inferred from a single simulation experiment. In all simulations we used a range of mutation rates, *μ,* values in combination with different sets of *P_sym_* values and bottleneck parameters that yielded substitution rate estimates fitting the observed substitution rates (see Figure 2A). The observed substitution rate was estimated as an average number of substitutions in neutral positions (Supplementary Note S1) that differs samples between two selected phylogenetic branches divided by two and normalized on the number of neutral positions and the divergence time of the corresponding branches in accordance with (Yu, Khachaturyan, et al. 2023). Similarly, for the plastid evolution simulation with an active partitioning mechanism, we used different *E_part_* values for different bottleneck regimes fitting the expected heteroplasmic sites estimates to the observed number of heteroplasmic sites. The *stratification scheme* parameter defines a set of probabilities of a particular cell to replace any other stem cell when dividing symmetrically. The scheme was designed in a manner that the cells could substitute only the neighbouring cells with cells at the end of the rows considered as neighbours (periodic boundaries). Thus, within one layer, all cells had equal contribution to mutation propagation.

To estimate the replicon copy number (*n*) parameter we analyzed 24 meristematic region samples of *Z. marina* ramets that belong to a single clone (Yu et al. 2020). The short-read data was aligned to the plastid and mitochondrial genomes by BWA-MEM with default parameters and the coverage was calculated directly by SAMtools. The average coverage was calculated for non-shared organelle genome areas and further normalized to the target coverage of the sequencing of 81x per diploid genome taken from Yu et al. 2020. The result replicon copy number per cell for the mDNA varied between 11-73 with an average of 40 copies per cell. The copy number of ptDNA ranged between 153-316 with an average of 216 copies per cell (Supplementary Table S1).

## Acknowledgements

We thank Hildegard Uecker, Agata Burian, Jessie Renton, Lei Yu, Devani Romero Picazo, and Fabian Nies for fruitful discussions and critical comments on the manuscript. We thank Fenna Stücker for graphical illustrations and Sergey Polevoy for his advice in algorithm implementation. MK was funded by the Helmholtz School for Marine Data Science (MarDATA), Grant No. HIDSS-0005, and HFSP Grant No. RGP0011/2022 (to TD). MS and TD are funded by ERC grant pMolEvol (Grant No. 101043835).

## Author contributions

MK, MS, TBHR, and TD conceived the study. MK and MS designed the modelling approach, MK performed the data analysis and developed the computer simulation. MK, MS, TBHR, and TD interpreted the results and wrote the manuscript.

