## Supplementary materials for "Heteroplasmy is rare in plant mitochondria compared to plastids despite similar mutation rates"

#### Supplementary information

### Supplementary Notes

**Supplementary Note S1: Detailed criteria for neutral position selection and mutation identification.**

In order to estimate the frequency of neutral fixed mutation and neutral heteroplasmic sites we first identified neutral positions, i.e., positions in which a detected mutation can be classified as neutral. Note that there are more neutral positions for fixed mutations than for heteroplasmic sites. An mtDNA positions was considered neutral for fixed mutations if the position was located in (i) non-coding areas; (ii) non-mtptDNA areas; (iii) non-microsatellite areas. Microsatellite loci were identified by searching the longest sequence of repeating (identical) nucleotide in the window of ten nucleotides upstream and downstream; such positions were retained only if the corresponding longest sequence was less than nine base pairs. An mtDNA position was considered neutral for heteroplasmic sites if (i) it fitted the criteria for neutral positions for fixed mutations; (ii) it belonged to non-NUMT areas; (iii) it had a coverage of at least 150x; (iv) it had a relative coverage between 0.3 and 1.5; (v) the median coverage of the whole mitochondrial genome was 50x or higher (that led to the exclusion of nine samples). Due to higher ptDNA copy number in comparison to mtDNA, we could apply less conservative filtration parameters for defining ptDNA as neutral. A ptDNA position was considered neutral for fixed mutations if it was located in (i) non-coding areas and (ii) non-microsatellite areas. A ptDNA position was considered neutral for heteroplasmic sites if it (i) fitted the criteria for neutral positions for fixed mutations; (ii) located in non-NUPT and non-mtptDNA areas; (iii) located in plastid single copy regions (SSC or LSC). We excluded plastid large inverted repeats (IRa and IRb) from the analysis as they are prone to frequent homologous recombination and therefore mutate at different rate than the single copy regions.

We identified neutral fixed and heteroplasmic variants in corresponding neutral positions using the following criteria. A single nucleotide polymorphism (SNP) in mtDNA was identified as a fixed mutations if (i) the position was classified as neutral and (ii) the variant allele frequency (AF) was $\geq$0.75. An SNP in mtDNA was identified as a heteroplasmic site if (i) the position was classified as neutral and (ii) the AF was between 0.05 and 0.95; here the AF was calculated from high quality reads supporting the variant or the wild type alleles only according to BCFtools mpileup v1.8 variant call results (Danecek et al. 2021). We additionally excluded (i) positions that were clustered together, (ii) positions for which the heteroplasmy can be explained by extra coverage of reads originated from another DNA region in at least one sample, and (iii) variants caused by differences in imperfect repeats. Among the three detected heteroplasmic sites, the variant in position 74138 was detected as fixed in most Californian samples and is therefore unlikely to be a genuine segregating *de novo* mutation.

We hypothesize that this variant arose due to biparental inheritance of mitochondria. Among the remaining two variable sites, position 124388 with AF=0.51 coming from one of the Mediterranean France (FR06 sample) population is not necessarily a genuine *de novo* mutation as well. This suspicion is raised due to other heteroplasmic position examples excluded during the filtration steps that grouped together Pacific populations and one of France samples (i.e., the position was variable in those populations). The AF of positions grouping Pacific populations together with the FR06 samples was between 0.46-0.47 that is similar in magnitude to AF=0.51 for position 124388. Thereby we conclude that the neutral heteroplasmic positions in the mitochondrial genome are 27884 (JS03, AF=0.91) and possibly also 124388 (FR06, AF=0.51). An SNP in ptDNA was identified as a fixed mutation if (i) the position was classified as neutral, (ii) the variant allele frequency (AF) was ≥0.75, and the relative coverage was ≥0.3. An SNP in ptDNA was identified as a heteroplasmic site if (i) the position was classified as neutral and (ii) the AF was between 0.05 and 0.95; here the AF was calculated from high quality reads supporting the variant or the wild type alleles only according to BCFtools mpileup v1.8 variant call results (Danecek et al. 2021). We additionally excluded the position 119234 due to a possible relation to a microsatellite of seven repeated adenines. Thereby the analysis resulted in six neutral heteroplasmic positions in the plastid genome (Table 1).

#### Supplementary Tables

**Supplementary Table S1: Replicon copy number in *Z. marina* Finish clone samples**. The sampled clone modules are named in accordance with Yu et al. (2020). The replicon copy numbers are estimated for the nucleus (ncDNA/cell), plastids (ptDNA/cell), and mitochondria (mtDNA/cell).

| **Module** | **ncDNA/cell** | **ptDNA/cell** | **mtDNA/cell** |
|---|---|---|---|
| M1 | 2 | 239 | 54 |
| M2 | 2 | 153 | 11 |
| M3 | 2 | 198 | 18 |
| M4 | 2 | 285 | 47 |
| M5 | 2 | 153 | 37 |
| M6 | 2 | 182 | 20 |
| M7 | 2 | 227 | 39 |
| M8 | 2 | 218 | 40 |
| M9 | 2 | 301 | 63 |
| M10 | 2 | 183 | 33 |
| M11 | 2 | 215 | 21 |
| M12 | 2 | 180 | 20 |
| M13 | 2 | 224 | 46 |
| M14 | 2 | 202 | 26 |
| M15 | 2 | 194 | 36 |
| M16 | 2 | 316 | 60 |
| M17 | 2 | 240 | 35 |
| M18 | 2 | 202 | 33 |
| M19 | 2 | 229 | 52 |
| M20 | 2 | 215 | 60 |
| M21 | 2 | 241 | 55 |
| M22 | 2 | 181 | 56 |
| M23 | 2 | 202 | 73 |
| M24 | 2 | 207 | 31 |

#### Supplementary Figures

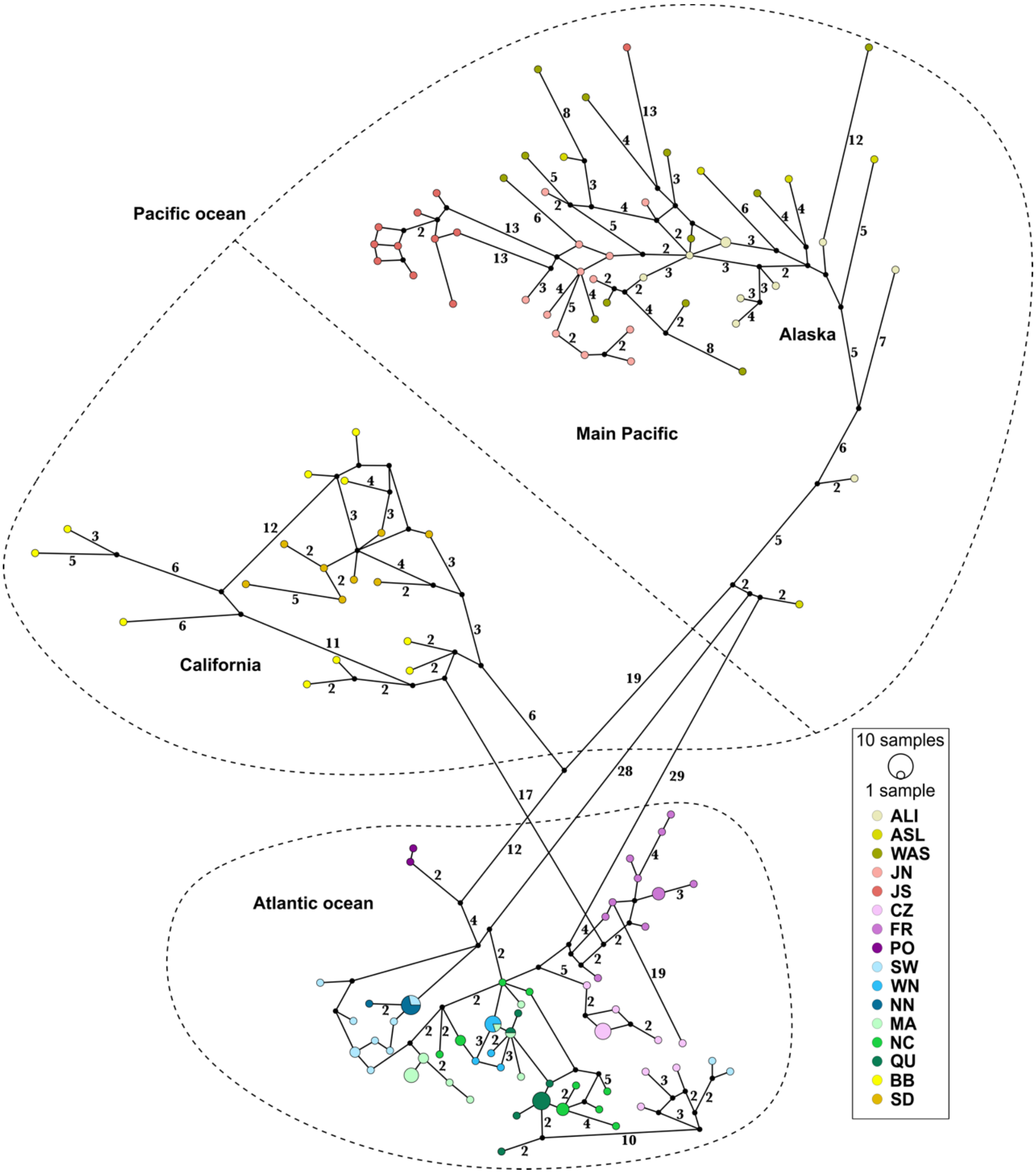

**Supplementary Fig. S1: Haplotype network of *Z. marina* mitochondrial genomes.** The haplotype network is reconstructed based on SNPs at neutral positions via the TCS Network method implemented in POPART v1.7 with default parameters (Clement et al. 2002; Leigh and Bryant 2015). The haplotypes are colored by the eelgrass population, split-colored circles indicate that a particular haplotype is shared between eelgrass populations, the size of the circle reflects the number of samples. Numbers on the edges show the number of mutation steps if more than one. Main groups of eelgrass populations are marked in accordance with the *Z. marina* phylogeny (Yu et al. 2023). For the population geographic locations see Yu et al. (2023) Figure 1. Population abbreviations: California: San Diego, California (SD), Bodega Bay, California (BB); Main Pacific: Washington state (WAS), Japan-North (JN), Japan-South (JS); Main Pacific (Alaska): Alaska-Izembek (ALI), Alaska-Safety Lagoon (ASL); Atlantic ocean: North Carolina (NC), Massachusetts (MA), Quebec (QU), Northern Norway (NN), Sweden (SW), Wales North (WN), Portugal (PO), Mediterranean France (FR), Croatia (CZ).

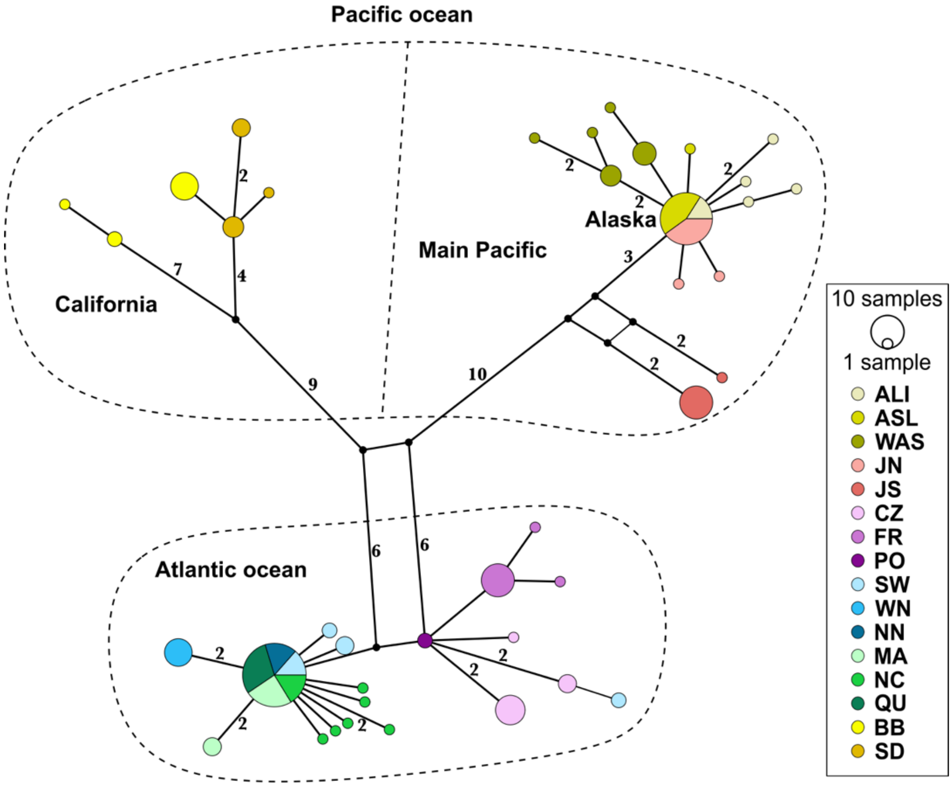

**Supplementary Fig. S2: Haplotype network of *Z. marina* plastid genomes.** The haplotype network is reconstructed based on SNPs at neutral positions via the Integer Neighbour-Joining method implemented in POPART v1.7 with default parameters (Leigh and Bryant 2015). The haplotypes are colored by the eelgrass population, split-colored circles indicate that a particular haplotype is shared between eelgrass populations, the size of the circle reflects the number of samples. Numbers on the edges show the number of mutation steps if more than one. Main groups of eelgrass populations are marked in accordance with the *Z. marina* phylogeny (Yu et al. 2023). The population abbreviations are the same as on the Supplementary Figure S1.

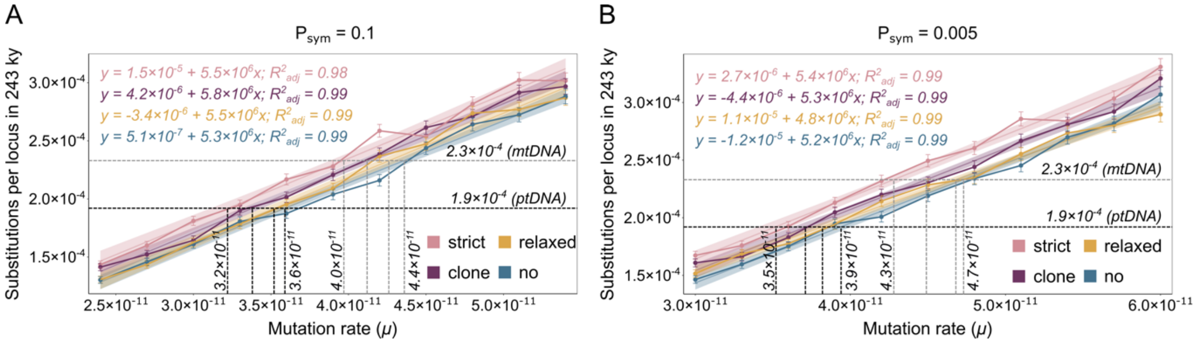

**Supplementary Fig. S3: Effect of the proportion of symmetric cell divisions ($P_{sym}$) on the mutation rate ($\mu$).** The number of mitochondrial substitutions per locus in 243,300 years in the *in-silico* experiments with $P_{sym} = 0.1$ (A) and $P_{sym} = 0.005$ (B) for different mutation rates ($\mu$). The dashed grey (mitochondrial genome) and black (plastid genome) lines indicate the $\mu$ values matching the observed number of accumulated fixed mutations. The color corresponds to the bottleneck regime, as in Figure 2A legend.

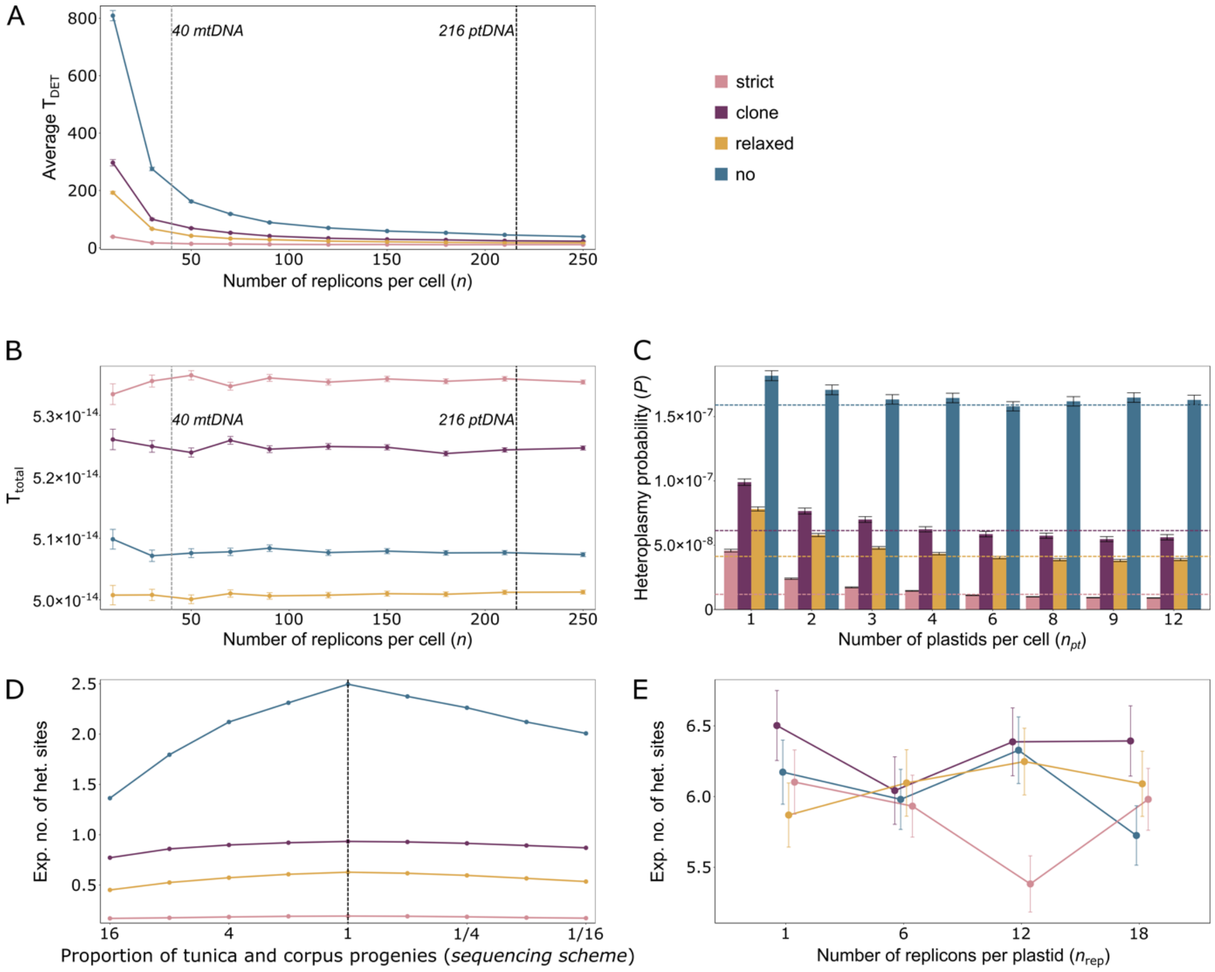

**Supplementary Fig. S4: Effects of individual model parameters**. a, The average detectable heteroplasmy time in cell divisions ($T_{DET}$) for different mitochondria-like replicon copy numbers per cell, i.e., with no additional population structure and random segregation. b, The total time in cell divisions ($T_{Total}$) in the same simulations as in (A) with the constant expected number of fixed mutations $M_{fix} = 500$. The vertical dashed lines correspond to the copy numbers of mtDNA (grey) and ptDNA (black) per cell. c, Effect of the population structure. The plot depicts the heteroplasmy probability estimated for different combination or ptDNA per plastid ($n_{rep}$) and plastids per cell ($n_{pt}$) numbers given a constant number of ptDNA copies per cell ($n = 216$), the mutation rates ($\mu$) that correspond to the estimates for the plastid genome (Figure 2A). The dashed lines reflect the mitochondrial heteroplasmy probability *P* ($n = 40$, mitochondrial $\mu$) calculated for the same simulation parameter set colored by the bottleneck regime. d, Effect of the *sequencing scheme* – the contribution of different apical initials to the sampled tissue. The simulation experiment is conducted for the mitochondrial parameter set. The proportion of progenies is assumed to be equal for stem cells of the same layer, while the proportion of L1 (tunica) layer progenies to L2 (corpus) progenies changes from 16 to 1/16. The expected number of heteroplasmic sites is estimated for 163 *Z. marina* samples. The dashed line reflects the equal number of progenies for all 20 stem cells that was used in other simulations as a default parameter. e, Effect of the number of plastid genome copies per plastid. The number of plastids per cell is $n_{pt} = 12$ in all experiments. The partitioning error ($E_{part}$) is set in accordance with the bottleneck regimes. The expected number of heteroplasmic sites is estimated for 163 *Z. marina* samples.

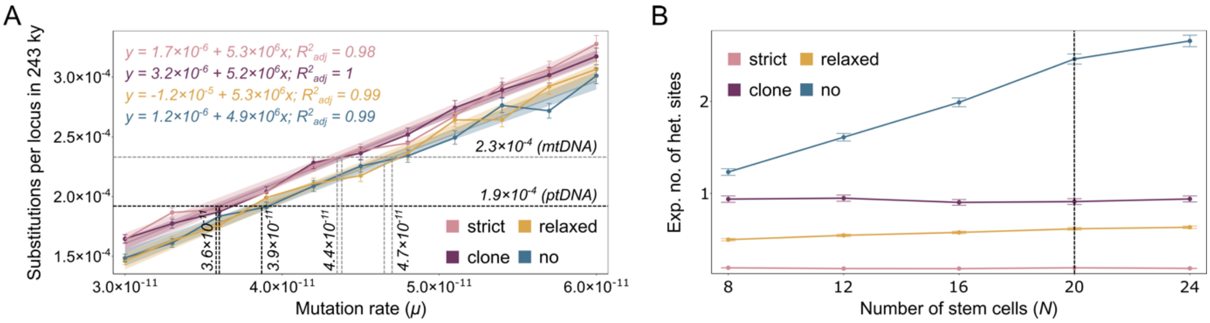

**Supplementary Fig. S5: Effect of the number of stem cells (*N*) on the expected number of heteroplasmic sites.** a, The number of fixed mitochondrial mutations per base pair in 243,300 years in the *in-silico* experiments for different mutation rates (*μ*), proportion of symmetric cell divisions $P_{sym} = 0.01$, $N = 8$ stem cells (four on the tunica layer and four on the corpus layer). The dashed grey (mitochondrial genome) and black (plastid genome) lines indicate the *μ* values matching the observed number of accumulated fixed mutations. The estimated *μ* values are similar to those estimated for $N = 20$ stem cells (Figure 2A). b, Expected number of heteroplasmic sites in 163 *Z. marina* samples estimated for different number of stem cells (*N*). In each simulation the stem cells were equally distributed between four rows – two in tunica layer and two in corpus layer. The vertical dashed line corresponds to the default number of stem cells in other simulation experiments.
